# AI-aided design of novel targeted covalent inhibitors against SARS-CoV-2

**DOI:** 10.1101/2020.03.03.972133

**Authors:** Bowen Tang, Fengming He, Dongpeng Liu, Meijuan Fang, Zhen Wu, Dong Xu

## Abstract

The focused drug repurposing of known approved drugs (such as lopinavir/ritonavir) has been reported failed for curing SARS-CoV-2 infected patients. It is urgent to generate new chemical entities against this virus. As a key enzyme in the life-cycle of coronavirus, the 3C-like main protease (3CL^pro^ or M^pro^) is the most attractive for antiviral drug design. Based on a recently solved structure (PDB ID: 6LU7), we developed a novel advanced deep Q-learning network with the fragment-based drug design (ADQN-FBDD) for generating potential lead compounds targeting SARS-CoV-2 3CL^pro^. We obtained a series of derivatives from those lead compounds by our structure-based optimization policy (SBOP). All the 47 lead compounds directly from our AI-model and related derivatives based on SBOP are accessible in our molecular library at https://github.com/tbwxmu/2019-nCov. These compounds can be used as potential candidates for researchers in their development of drugs against SARS-CoV-2.

## Introduction

The emerging coronavirus SARS-CoV-2 has caused an outbreak of coronavirus disease (COVID-19) worldwide.^1^ As of March 2, 2020, more than 90,000 people have been infected by SARS-CoV-2 and more than 3000 people have been reported dead according to Johns Hopkins Coronavirus map tracker.^2^ The numbers of infection and death are still increasing. To face the considerable threat of SARS-CoV-2, it is urgent to develop new inhibitors or drugs against this deadly virus. Unfortunately, since the outbreak of severe acute respiratory syndrome (SARS) eighteen years ago, there has been no approved treatment against the SARS coronavirus (SARS-CoV),^3^ which is similar to SARS-CoV-2. Repurposing potential drugs such as lopinavir and ritonavir also failed to SARS-CoV-2 injected patients.^4^ Structure-based antiviral drug design with a new artificial intelligence algorithm may represent a more helpful approach to get the SARS-CoV-2 targeted inhibitors or drugs. Thanks to the prompt efforts of many researchers, we have several pieces of important information about this vital virus genome and protein structures. We now know that the non-structural protein 5 (Nsp5) is the main protease (M^pro^) of SARS-CoV-2 and it is a cysteine protease, which also been called “3C-like protease” (3CL^pro^). Moreover, we know that the 3D structure of 3CL^pro^ is very similar to SARS-CoV with a sequence identity of >96% and 3D structure superposition RMSD_Cα_ of 0.44 Å as shown in Figures S1 and S2.

3CL^pro^ has been reported as an attractive target for developing anti-coronaviral drugs: 1) this protease is highly conserved in both sequences and 3D structures;^5^ 2) 3CL^pro^ is a key enzyme for related virus (including SARS and SARS-CoV-2) replication; 3) it only exists in the virus, not in humans. Developing specific antiviral drugs targeting 3CL^pro^ of the specific virus has shown significant success; for example, both approved drugs lopinavir and ritonavir can completely occupy the substrate-binding site of 3CL^pro^ to break down the replication of human immunodeficiency virus (HIV). However, due to the large difference between HIV and SARS-CoV-2 3CL^pro^, lopinavir and ritonavir were validated ineffective for inhibiting SARS-Cov-2.^4^ On the other hand, the substrate-binding site of 3CL^pro^ is almost the same between the SARS-CoV-2 and SARS as Figure S3 presents. The developed potential inhibitors and drug-design experience targeting SARS-3CL^pro^ may also be applicable to SARS-CoV-2. For example, the recently solved structure of SARS-CoV-2 3CL^pro^ (PDB ID: 6LU7) indicates that the developed inhibitor N3,^6^ which is a covalent inhibitor derived from non-covalent inhibitors against SARS can also bind SARS-CoV-2 3CL^pro^ with a similar binding conformation (Figure S4).

All the above available information paved a way to design new targeted covalent inhibitors (TCI)^7^ against SARS-CoV-2. A successful TCI against 3CL^pro^ must first be able to fit in the binding site of 3CL^pro^ with an appropriate pose that keeps its reactive groups close enough to the Cys145, which then undergoes a chemical step (nucleophilic attack by Cys145) leading to the formation of a stable covalent bond as presented in the scheme below:

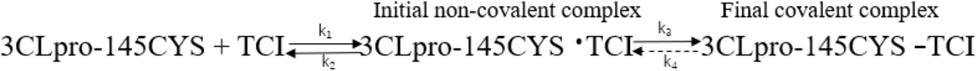

TCIs usually have a longer target residence time than the relative non-covalent inhibitors in theory given the following: 1) For the inhibition, k_1_ must be larger than k_2_, and thus the non-covalent binding is determined by the equilibrium constant k_1_/k_2_; 2) TCIs have the chemical reaction step with the target, where usually k_3_ ≫ k_4_; and 3) for TCIs, the binding process is controlled by k_1_ k_3_/(k_2_ k_4_), which is bigger than k_1_/k_2_ in non-covalent inhibitors. In some extreme cases k_4_ = 0, and hence, these irreversible inhibitors covalently bind the target until the target disappears.^7, 8^

Considering the inhibitors of SARS 3CL^pro^ may be also bio-active to SARS-CoV-2, we have created a molecular library including all the reported SARS-3CL^pro^ inhibitors (284 molecules)^9, 10, 11, 12, 13, 14, 15, 16, 17, 18, 19, 20, 21, 22, 23, 24, 25 26, 27, 28, 29^ and we will also add new validated molecular structures into this library with our research progresses. To date, there are no clinically approved vaccines or drugs specifically targeting SARS-CoV-2. Thus, with the hope to discover novel candidate drugs targeting SARS-CoV-2, we combine artificial intelligence (AI) with the structure-based drug design (SBDD) to accelerate generating potential lead compounds and design TCIs.

AI, especially deep learning, has been applied in predicting molecular properties^30, 31, 32, 33^ and designing novel molecules^34^. Unlike earlier deep-learning molecular design by adding single atom one at a time^34, 35^, our approach explores new molecules by adding a meaningful molecular fragment one by one, which is not only computationally more efficient but also chemically more reasonable. To make our AI model work well, the first step is to prepare the molecular fragment library as shown in Figure 1. We used our collected SARS-CoV 3CL^pro^ inhibitors (284 molecules) as the initial molecule database targeting SARS-CoV-2 3CL^pro^. Then, we split this set of molecules into fragments with a molecule weight no more than 200 daltons. Both of the collected inhibitors and the fragments are supplied in https://github.com/tbwxmu/2019-nCov. Then we applied an advanced deep Q-learning network with the fragment-based drug design (ADQN-FBDD) for generating potential lead compounds. Noted, if researchers have enough experience and internal lead compounds or biased fragments, they can inject all such information into ADQN-FBDD by manually adding the lead compounds and biased fragments to the corresponding files. By using the same fragments directly from existing bioactivate molecules, our ADQN-FBDD agent can easily access the potential chemical space focused on 3CL^pro^ of SARS-CoV-2.

**Figure 1.**
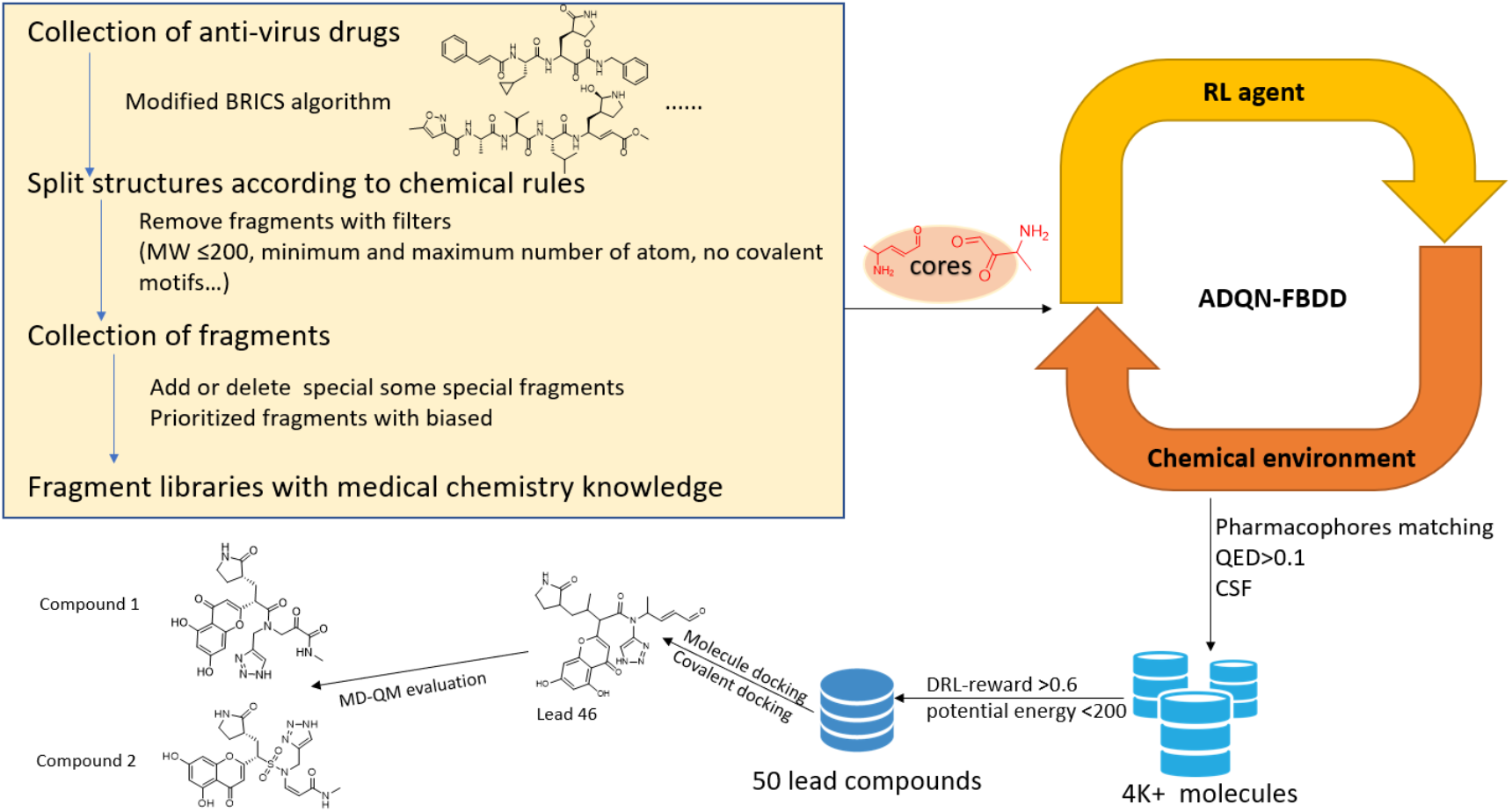
Flowchart for our SARS-CoV-2 3CL^pro^ lead compounds development.

After ADQN-FBDD automatically generated novel compounds targeting this new virus 3CL^pro^, we obtained a covalent lead compound library with 4,922 unique valid structures. In total, 47 of these compounds were selected with high scores from our AI-model’s reward function. Then these molecules were further evaluated by docking and covalent docking studies. The lead compound #**46** with a high covalent docking score attracts our attention, which also has a low difference between non-covalent and covalent docking pose among the 47 lead compounds. After carefully checking the lead #**46**’s interaction mode with 3CL^pro^, we believe there is still much space to optimize lead it. Then we designed a series of derivatives from compound #**46** based on our chemical biology knowledge and the structure-based optimization policy. All the generated molecular structures are published in our code library https://github.com/tbwxmu/2019-nCov. We encourage researchers who are interested in finding a potential treatment for this viral infection to synthesize and evaluate some of these molecules for treating COVID-19.

## Results

Integrating the double dueling deep Q learning with fixed q-targets and prioritized experience replay enables our agent ADQN stable and efficient during learning from the chemical environment. Combining the state-of-the-art AI algorithm with the idea of FBDD as presented in Figure 2, ADQN-FBDD is flexible and efficient to access the focused chemical space targeting the SARS-CoV-2 3CL^pro^. Based on the configurations targeting the SARS-CoV-2 3CL^pro^, ADQN-FBDD generated a potential lead library containing 4,922 unique molecular structures (Supplementary dataset in https://github.com/tbwxmu/2019-nCov). To narrow our focus to a smaller set of molecules for analysis, we elaborately defined filter rules (QED>0.1 and DRL-reward>=0.6) and the detailed information of the rules can be found in the Methods section. And then 47 unique molecules (Table S1) were kept for the next non-covalent docking and covalent-docking evaluation. These 47 virtual leads display suitable 3D-complexity with common characteristics of peptidomimetics and protein-protein interaction (PPI) inhibitors. They are mainly ranked by covalent-docking scores, considering covalent-docking also contains the scores of non-covalent docking.^36^ And we also paid attention on the RMSD difference between the covalent binding and non-covalent binding poses based on all heavy atoms. Finally, we selected the lead molecule #**46** as displayed in Figure 3 with both a good covalent docking score and a small RMSD value (Table S1). We further optimized it and get a series of derivatives based on the SBDD approach.

**Figure 2.**
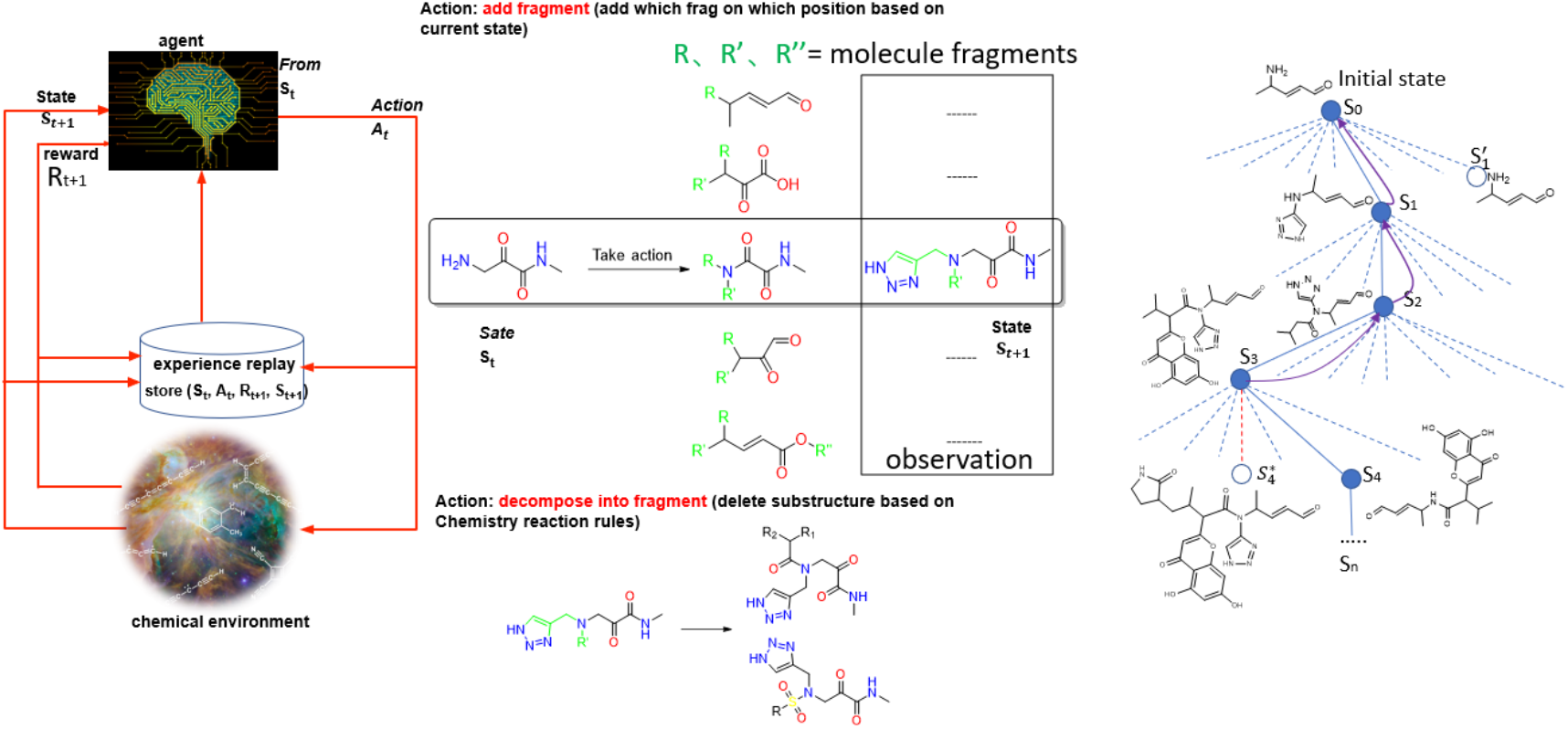
A: Framework of ADQN-FBDD. B: Examples for explaining the fragment-based actions. C: The solid lines represent taken actions including the addition/deletion of a different fragment or no modification during each step. The dashed lines represent actions that our reinforcement learning agent considered but did not make. After the first three actions to S_1_, S_2_, and S_3_, the fourth action to S_4_ was an exploratory action, meaning that it was taken even though another sibling action, the one colored in red dashed line leading to 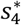, was ranked higher. Exploratory action does not result in any learning, but each of our other actions does, resulting in updates as suggested by the purple curved arrow in which estimated values are moved up the tree from later nodes to earlier ones.

**Figure 3.**
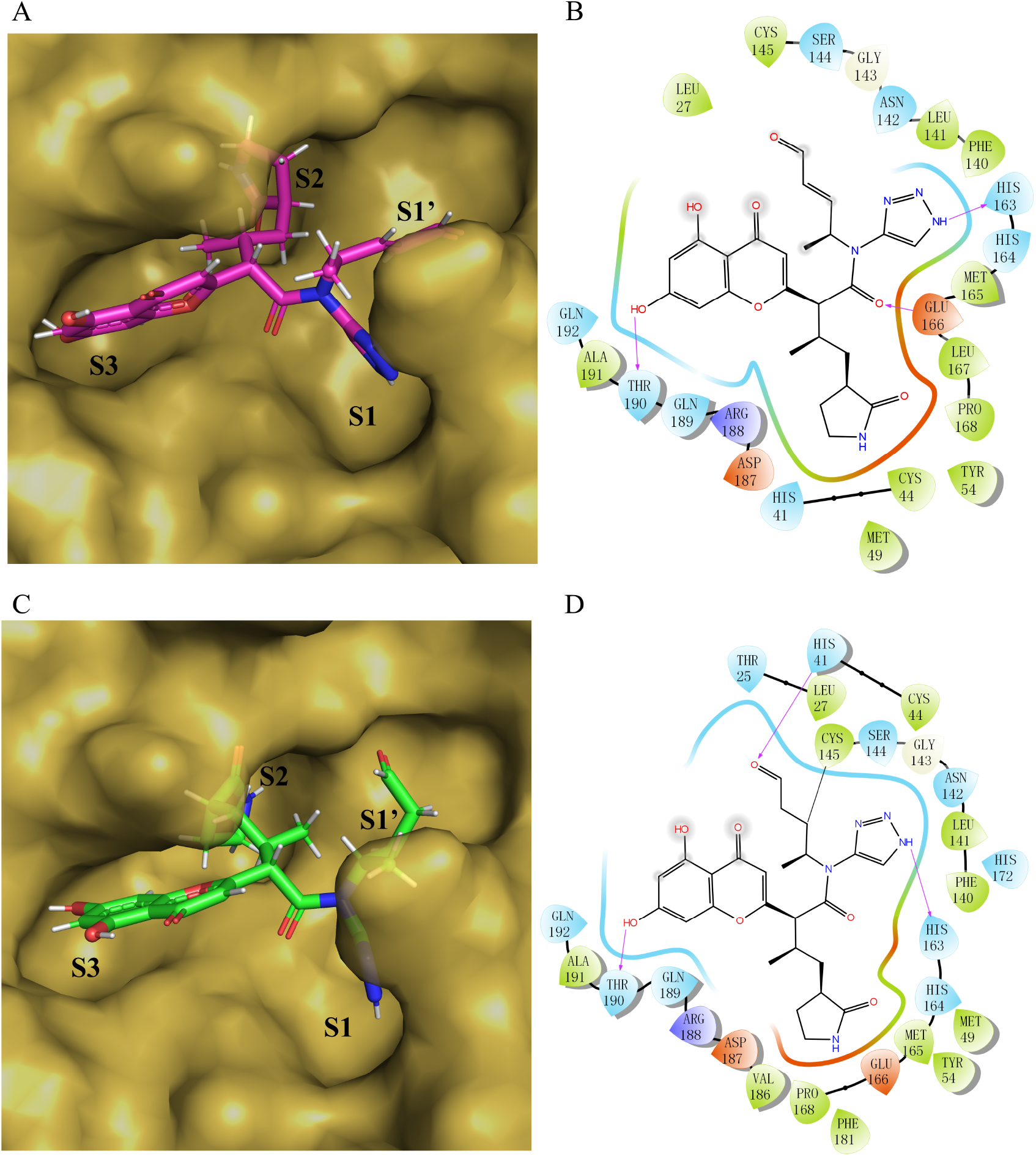
Lead compound #**46** generated by AI. A: Non-covalent molecular docking model of SARS-CoV-2 3CL^pro^ (brown surface) with the bound lead compound #**46 (**magenta sticks). The triazole ring binds to the S1 subsite of the catalytic active center, the covalent fragment of α, β-unsaturated aldehyde binds to S1’ subsite, β-lactam ring binds to S2 subsite and 5,7-dihydroxy chromone binds to S3 subsite. B: A two-dimensional (2D) view of non-bonding interaction of lead compound #**46** in complex with 3CL protease based on non-covalent docking. The triazole ring, i.e., keto-amide, phenolic hydroxy forms hydrogen bond (H-bond) with His163, Glu166, and Thr190, respectively. C: Covalent docking model between compound #**46 (**green sticks) and 3CL protease (brown surface) exhibits a similar docking model as non-covalent docking. D: A 2D view of ligand interaction between compound #**46** and protease under covalent docking. The triazole ring forms an H-bond with His163; the fragment of α, β-unsaturated aldehyde forms a covalent bond with Cys145, i.e., the key residue in the catalytic center of the protease, resulting in covalent inhibition; aldehyde carbonyl forms an H-bond with His41; and besides, the hydroxyl of chromone at position 7 forms an H-bond with Thr190.

Molecule #**46** was ranked number 1 based on the covalent docking score and its interaction model with the binding site was carefully checked as shown in Figure 3. Although compound #**46** has the best covalent docking score, it has the alerting group aldehyde. Considering there is still much space for compound #**46** to fill in the S1’ subsite and α-ketoamides may be good to fit the oxyanion hole (Figure 4A) of 3CL^pro^,^3^ we replaced the aldehyde by formamide and also replaced the 1,4 Michael acceptors by alpha-ketoamides. Thus, we optimized compound #**46** to compound **46-14-1**(Figure 4 A and B).

**Figure 4.**
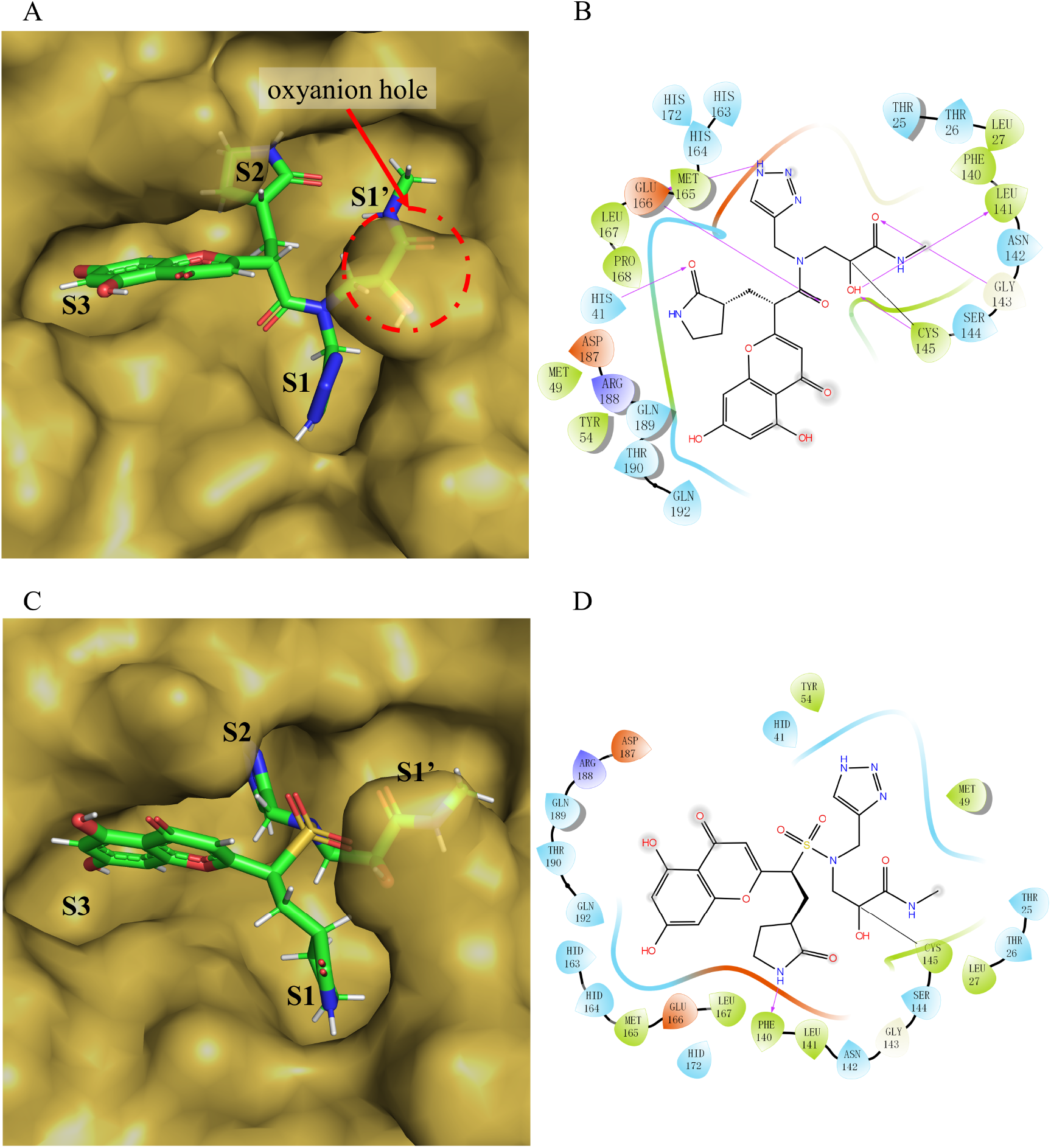
The covalent binding models of compound **46-14-1** and **46-14-2** in complex with the SARS-CoV-2 3CL^pro^. A: The covalent docking model between compound **46-14-1** (green sticks) with 3CL protease (yellow-orange surface). The oxyanion hole formed by the segment of α-ketoamides is shown in the red circle. B: The detailed view of the interactions between the compound **46-14-1** and 3CL^pro^. C: The covalent docking model between compound **46-14-2** with 3CL protease. The molecule **46-14-2** is shown as green sticks and the protein is shown as brown surface. D: The 2D view of the interactions between the compound **46-14-1** and 3CL^pro^.

The non-bonding interaction between compound **46-14-1** and SARS-CoV-2 3CL^pro^ is mainly hydrogen bond. The carbonyl group of covalent scaffold α-ketoamide forms H-bonds with Leu141 and Gly143 as a hydrogen acceptor or a hydrogen donor. The hydrogen on the nitrogen atom of the triazole ring forms an H-bond with Glu166, while Glu166 also forms an H-bond with the carbonyl group on the main chain. The oxygen in the β-lactam ring forms an H-bond with His41. In order to enhance the polarity of the compounds, sulfonic groups were introduced to replace the ketone carbonyl groups on the main chain and sulfonamides **46-14-2** were obtained. The covalent docking model of compound **46-14-2** with SARS-CoV-2 3CL^pro^ is shown in Figure 4 C and D.

In order to make the compound **46-14-2** fit the active pocket with a higher affinity, we added a carbon atom to the sulfonic acid group of the original molecule, extending the carbon chain to increase the flexibility of the molecule, and obtained another optimized compound **46-14-3**. The mode of covalent docking with SARS-CoV-2 3CL^pro^ is shown in Figure 5. Due to the introduction of carbon atoms and the enhancement of molecular flexibility, the β-lactam ring can be inserted deeper into the S2 pocket and the other fragments of the compound can better adapt to the S1, S1’ and S3 subsites. The α-carbonyl carbon on the α-ketoamide of compound **46-14-3** forms a covalent bond with the key residue Cys145 on the protease, but the main non-bond interaction is still hydrogen bond (indicated by a yellow dash). The triazole ring mainly forms H-bonds with Phe140 and Glu166 residues in the S1 pocket; α-ketoamide covalent binding fragment mainly forms H-bonds with key amino acid residues Cys145, Gly143 and Ser144 in the S1’ subsite, which forms an oxyanion hole in the red circle in Figure 9; the β-lactam side chain mainly forms H-bonds with residues Tyr54 and AsS187 in the S2 pocket; the chromone scaffold mainly forms H-bonds with key residues Thr190 and Gln192 in the S3 pocket. In addition, the oxygen on the sulfonyl group of the main chain forms an H-bond with Glu166 in the S1 pocket.

**Figure 5.**
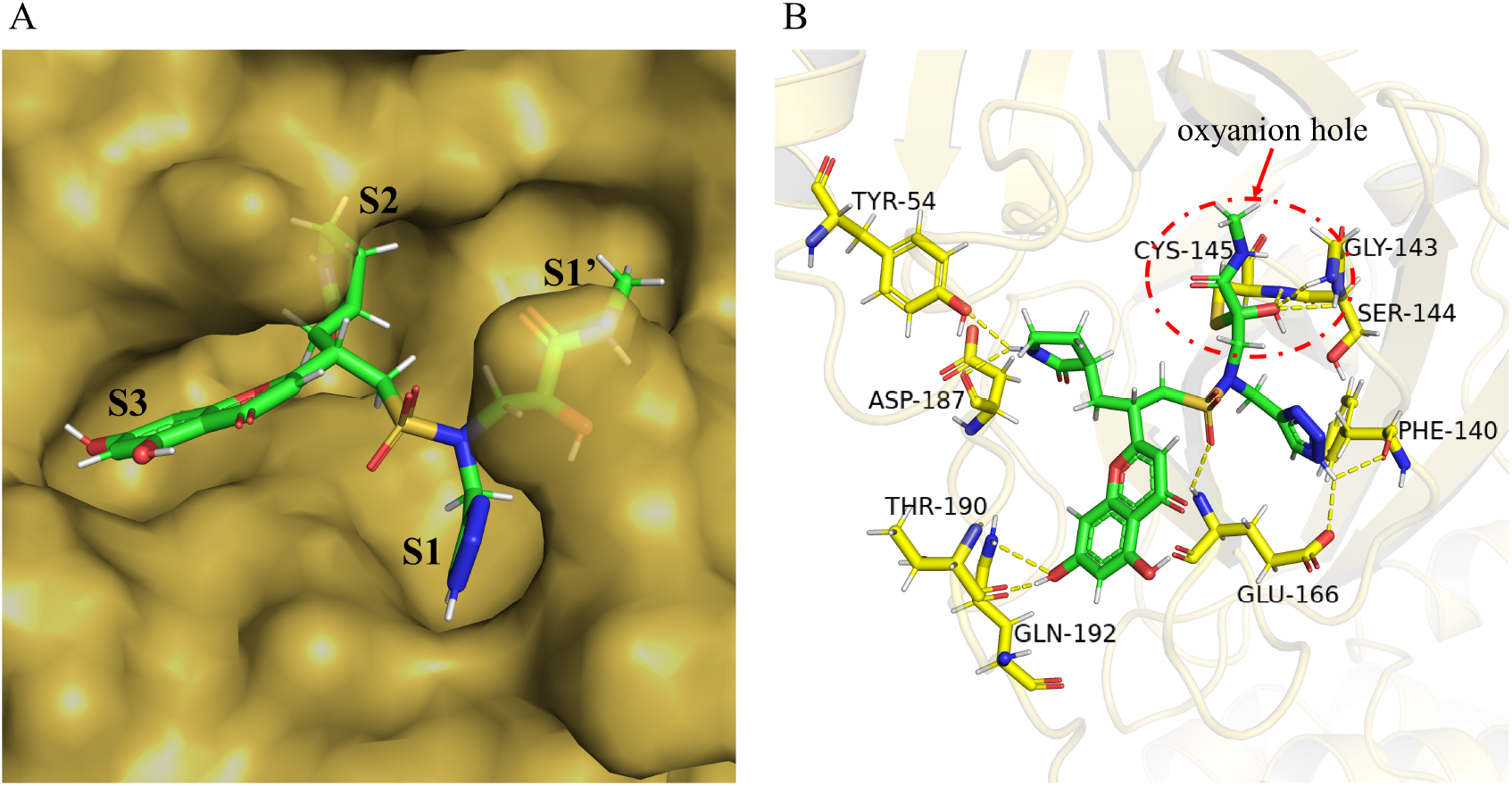
The covalent binding model of Compound **46-14-3** in SARS-CoV-2 3CL^pro^. A: Surface representation of SARS-CoV-2 3CL^pro^ (brown) complexed with **46-14-3** (green sticks). B: A stereo view showing **43-14-3** bound into the substrate-binding pocket of the SARS-CoV-2 3CL^pro^ at 4 Å. The molecule **43-14-3** is shown as green sticks. Residues forming the H-bond are shown as yellow sticks. And the yellow dashes represent the H-bonds, as well as the oxyanion hole, is in the red circle region.

The representatives were selected (**46-14-1**, **46-14-2** and **46-14-3** as Figure 6 displays), which may be further evaluated by molecular dynamics simulation to get the binding free energy and by quantum chemical calculation to get the reaction energy barrier. Meanwhile, **46-14-1**, **46-14-2** and **46-14-3** are chosen as our candidates for chemical synthesis and anti-SARS-CoV-2 activity testing, which is ongoing.

**Figure 6.**
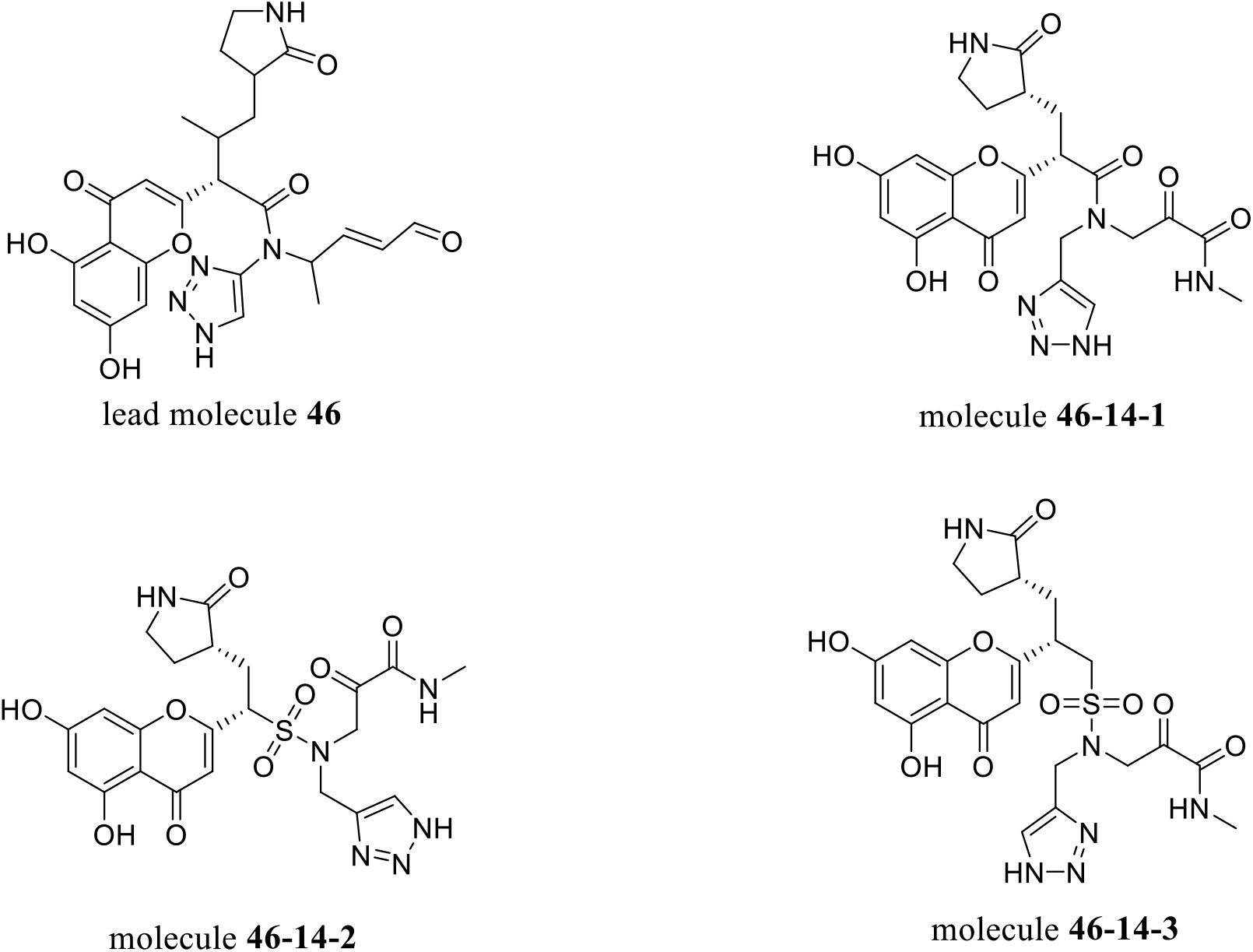
Structures of optimized compounds.

## Discussions

Computational approaches are particularly important for emerging diseases given the urgent need to provide timely solutions. In this work, our robust and efficient computational method and pipeline for designing compounds can provide useful drug candidates for treating SARS-CoV-2 infections. For more information about our AI model generated leads and SBDD optimized derivatives, please go to the library https://github.com/tbwxmu/2019-nCov. These candidates or their variants have a good chance to produce valid leads of anti-COVID-19 drugs. It is understood that the computational design requires experimental validations. While we are exploring experimental validations ourselves, we like to release these candidates promptly for other researchers to accelerate the development of anti-COVID-19 drugs given the emergency of seeking treatments for the disease.

Comparing with other deep RL methods, ADQN-FBDD has several highlights: 1) It directly modifies and generate molecular structures without format conversion problems while some other tools such as Insilico Medicine’s GENTRL (https://insilico.com) may generate invalid SMILES output. 2) Most of the generative models require pre-training on a specific dataset and then produce the molecules with high similarities to a given training set. For example, the best molecule from Insilico Medicine for their target DDR1 was actually very similar to the kinase inhibitor Iclusig (ponatinib) on the market.^37^ ADQN-FBDD does not need pre-training at all and has a capacity to generate novel molecules. 3) The process of generating molecules is very efficient and effective as ADQN-FBBD is molecular fragment-based growing with the knowledge of chemical reactions, while other models are all atom-based with no rules of chemical reactions at all^34, 38, 39^. 4) ADQN-FBDD is highly flexible and user-friendly for the medicinal chemists, who can easily inject their drug discovery experience into the reward function to guide the novel molecule generation. Our ADQN-FBDD and related pipeline can be used not only for designing anti-COVID-19 drugs but also other structure-based drug discoveries, especially for emerging infetious diseases that require treatments timely.

## Methods

### Markov Decision Process for Molecule Generation

Intuitively, the problem of chemical structure graph generation is formulated as learning a reinforced agent, which performs discrete actions of chemical reaction-based fragment addition or removal in a chemistry-aware Markov Decision Process (MDP). Formally, the components of MDP) include *M* = {*S, A, P, R, γ*}, where each term is defined as follows:

*S* = {*S*_*t*_} denotes the state space containing all possible intermediate and final generated molecular graphs. Each *s*_*t*_ is a tuple of (*s*, *t*). *s* stands for a valid molecule structure and *t* is the time step. For the initial state *s*_0_, the structure can be represented as a specific core as Figure 2C indicates or randomly chosen from the prepared fragment library at time *t* = 0. We also limit the maximum number of time steps *T* in our molecule fragment based MDP, which defines the set of terminal states as {*s*_*t*_|*t*=*T*} containing all the states with the number of steps reaching the maximum allowed value *T*.

Ac={*A*_*t*_} denotes a set of actions that describe the modification made on the current molecule structure at each time step *t*. Here, each action can be classified into three categories: fragment addition, fragment deletion, and no modification.

*P* =*p*(*s*_*t*+1_ |*s*_*t*_ … *s*_0_) =*p(s*_*t+1*_|*s*_*t*_, *a_t_)* is the basic assumption in MDP. The state transition probability, which specifies the next possible state given the current state and action at time step t. Here, we define the state transition to be deterministic. For example, as Figure 2C indicates S_0_ to S_1_, by adding a 1*H*-1,2,3-triazol-4-yl fragment on *S*_*0*_, the next state *S*_*1*_ will be the new structure of added 1*H*-1,2,3-triazol-4-yl with a probability of 1.

*R* is the reward function that specifies the reward after reaching state *S*_*t*_ *γ∊*(0,1] is the discount factor and typically γ = 0.9 in our study. In our framework, the state always has a valid and complete chemical structure at each step as Figure 2C indicates. A reward is given not just at the terminal states, but at each step. Both intermediate rewards and final rewards are used to guide the behavior of the reinforcement learning (RL) agent. So, there is no delayed or sparse reward issue as many other reinforced frameworks suffered.^40, 41^ Furthermore, to ensure that the last state is rewarded the most, we use *γ*^*T*−*t*^ to discount the value of the rewards at state *s*_*t*_. In addition, our reward function can directly integrate the experience of medicinal chemists. For example, given a core of interest, medicinal chemists may add biased fragments of their interests and their input can be used to design a reward function that gives high reward signal values to biased fragments so that ADQN-FBDD may have a better chance to generate the desired structures.

### Chemical Environment Design

In our RL framework, the chemical environment receives action *a*_*t*_ from the agent and emits scalar reward r_t_ and state 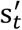 to the agent as Figure 2A shows. Note that the definition of the environment state is different from the general approach that the environment state is only the environment’s private representation invisible to the agent. We define the state of the chemical environment 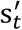 as the intermediate generated molecule structure at time step t, which is fully observable by the RL agent. Simply, the environment’s state 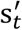 is the agent’s state *s*_*t*+1_. For the task of molecule generation, the environment incorporates rules of chemistry. In our study, chemistry rules are not only the basic chemical valency, but also the rules about adding and removing the fragments derived from known inhibitors based on chemical reactions. The detail information of 45 defined chemical reaction rules is presented in Table S2.

### Agent Design

As indicated in Figure 2A, the basic model of our ADQN-FBDD is an advanced Q-network. The goal of molecule generation is equally to fit a Q function *Q*(*s*_*t*_, *a*_*t*_) to make the agent choose the action *a*_*t*_ at state *s*_*t*_ that maximizes the future expected γ-discounted cumulative rewards with policy π. Mathematically, given the agent’s policy π, the value of the state-action pair Q^π^(*s*_*t*_, *a*_*t*_) and the value of state V^π^(*s*_*t*_) are defined as, respectively:

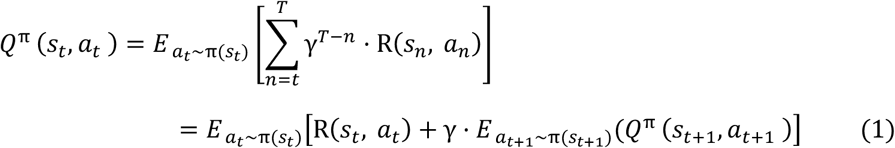

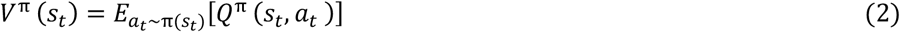

where 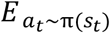 is the expectation within policy π on state *s*_*t*_ taken *a*_*t*_ and R(*s*_*n*_, *a*_*n*_) denotes the reward at step *n*. Value function Q^π^(*s*_*t*_, *a*_*t*_) measures the value of taking action *a*_*t*_ on state *s*_*t*_. V^π^(*s*_*t*_) is the value of being at state *s*_*t*_ means how good to be in this state. Obviously, V^π^(s_t_) can be seen as a part of Q^π^(*s*_*t*_, *a*_*t*_). Then, the rest part from Q^π^(*s*_*t*_, *a*_*t*_) can be defined as the so-called advantage function A^π^(*s*_*t*_, *a*_*t*_)^42^ as:

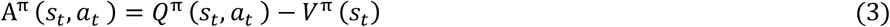

Intuitively, the advantage value shows how advantageous selecting the action is relative to the others at the same given state. Then Eqn. (2) can be rewritten according to Eqn. (4):

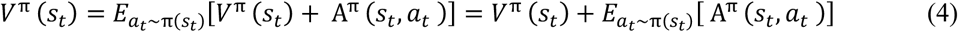

Obviously, 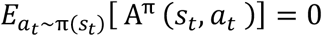. To avoid the issue of identifiability, we deduct the mean value from the prediction and the Q-function of dueling DQN can be defined as:

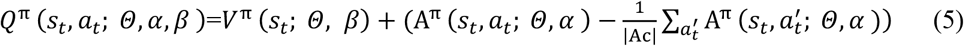

Note that *Θ, α* and *β* come from the dueling Q-network as Figure 7 indicates. |Ac| is the size of action space and 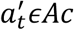. To make our RL agent more robust to be more stable learning and to handle the problem of the overestimation of Q-values, double Q-network^43^ and fixed Q-targets^44^ are also incorporated:

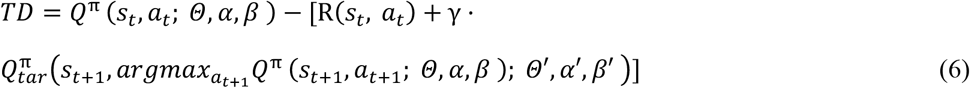

where, TD is the temporal-difference; 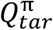 is another dueling DQN network as the target network and its parameters (*Θ′, α′, β′*) keep fixed and copy from the dueling DQN Q^π^ every *m* steps (*m*=20 as we used). To update the parameters (*Θ, α, β*) from the dueling DQN as Figure 7 displays, we can train our RL agent by minimizing the loss function:

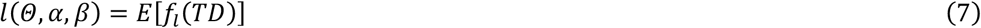

where ***E*** is the expectation. As the L2 loss has a disadvantage of the tendency to be dominated by outliers, we use the Huber loss as the loss function *f*_*l*_:

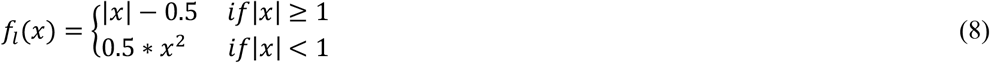

**Figure 7.**
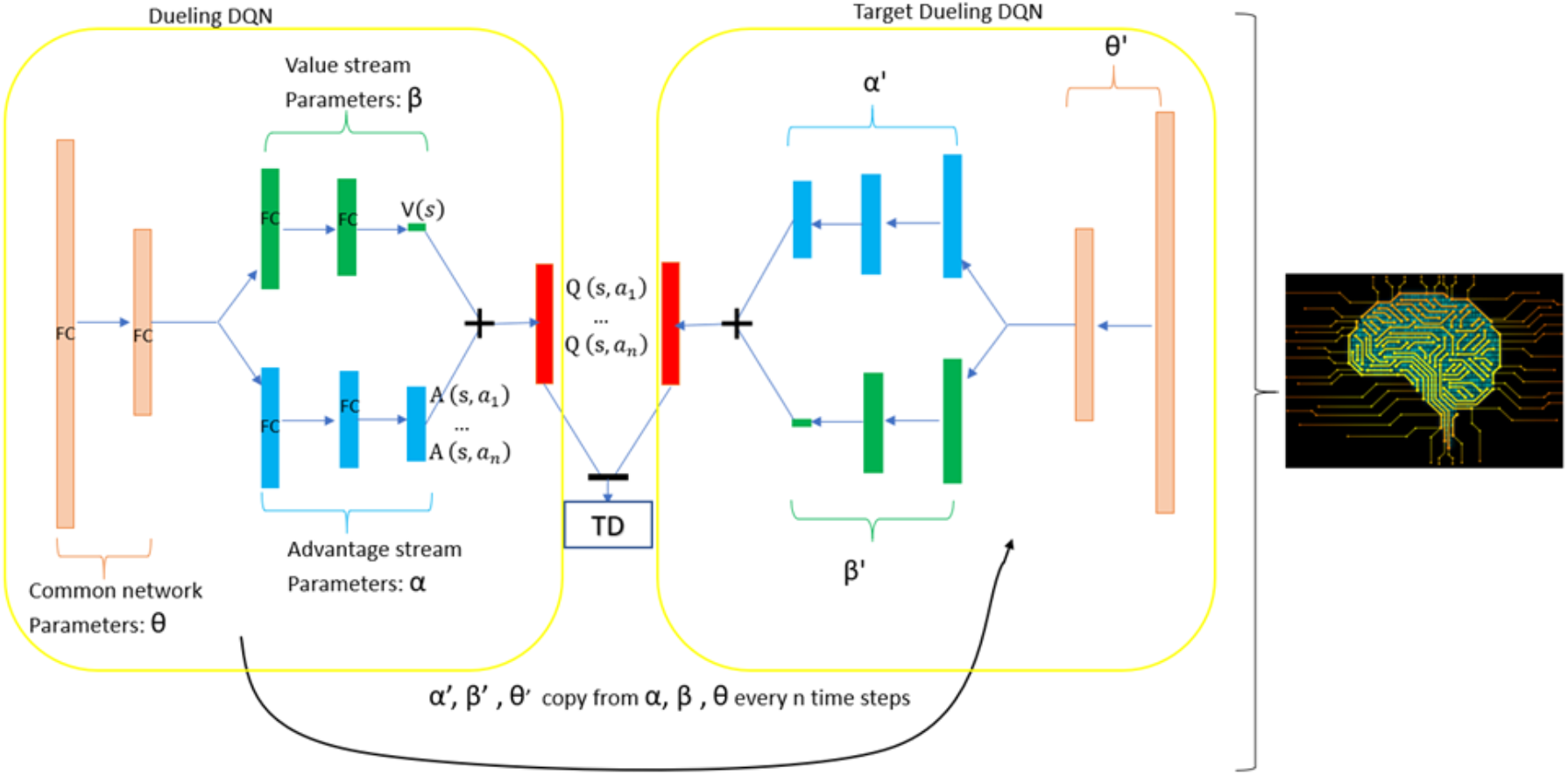
Architecture of used advance deep Q-learning network (ADQN). TD is the temporal difference error.

### Prioritized Experience Replay

Prioritized Experience Replay (PER)^45^ is a technique to enable reinforcement learning agents remember and reuse experience from the past, and to replay important transitions more frequently. PER is very useful for replaying some less frequent experiences. Here, we use the same code of “Prioritized Replay Buffer” from open AI’s gym with version 0.15.4.^46^ Finally, our RL agent is the double dueling deep Q learning with fixed q-targets and prioritized experience replay.

### Fragment Library Design

The fragment-based approach to drug discovery (FBDD) has been established as an efficient tool in the search for new drugs.^47^ The idea of FBDD is that proper optimization of each unique interaction in the binding site and subsequent incorporation into a single molecular entity should produce a compound with a binding affinity that is the sum of the individual interactions. However, the widely used fragment libraries only consider the diversity of fragments, such as the ZINC fragment database. They have a very low probability to exhibit the desired bioactivity for a given protein.

To combine the idea of FBDD with our RL framework, we first collected and built a SARS-CoV-2 3CL^pro^ inhibitor dataset containing 284 reported molecules. Then using the improved BRICS algorithm^48^ to split those molecules to get the fragment library target on SARS-CoV-2 3CL^pro^ as the flowchart displayed in Figure 1’s yellow box. An elaborate filtering cascade accompanied by manual inspection and the rules can be changed based on the needs in different studies. Our fragment library contains 316 fragments with molecular weight <200 daltons, the minimum number non-hydrogen atom >1 and the maximum ≤ 25. The fragments directly from the existing inhibitors based on the chemical retrosynthetic rules are always true substructures and may have a high quality of bioactivity targeting 3CL^pro^. It is worth noting that the quality of the designed fragment library directly affects the behavior of the chemical environment of ADQN-FBDD.

### Core Selection

Studies have identified various scaffolds or core structures that have privileged characteristics in terms of the activity of a certain target.^49, 50, 51^ Core structure selection is the starting point in a scaffold-based drug discovery. Choosing or designing a proper initial scaffold is never trivial, and medicinal chemists may need enough experience to get such a skill. Luckily, there are serval reported privileged core structures targeting SARS M^pro^.^3, 52^ Here, we chose 4-aminopent-2-enal and 3-amino-2-oxobutanal as the starting cores as Figure 8 displays, because both cores have been validated to generate covalent bonds with the Cys145 of SARS or SARS-CoV-2 3CL^pro^.

**Figure 8.**
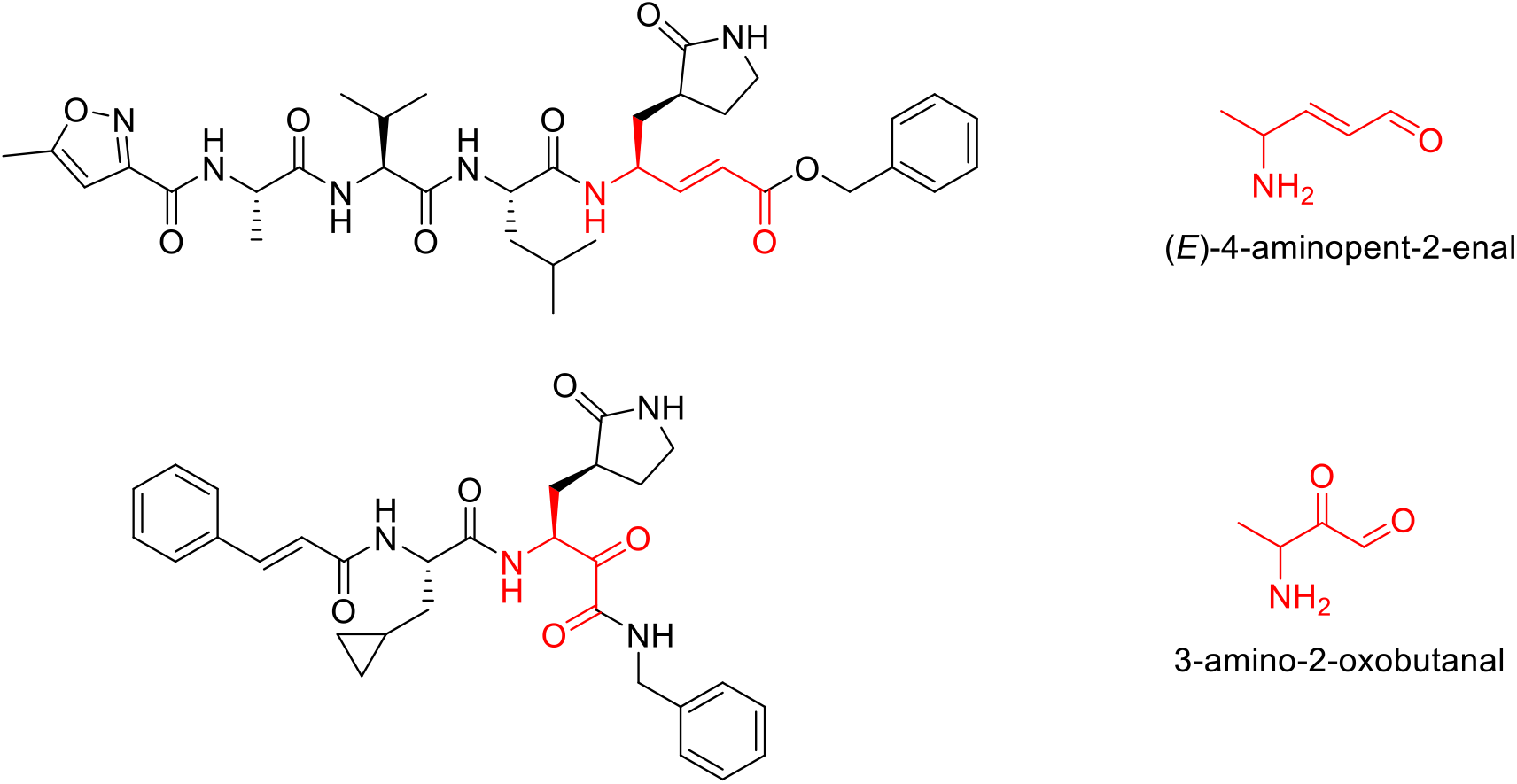
Structures of the chosen cores

### Reward Design

Most reported RL methods using the complete structure information of a positive drug or inhibitor as the template,^38, 39^ and they design a reward function for the RL agent to learn to regenerate the template structure or generate highly similar structures to the template. This way may be useful in testing the performance of RL methods but not suitable in a real-world drug design because no one knows the complete structural information of the novel molecule. A more practical approach is to learn the structure features of existing drugs or inhibitors to local, focused chemical space for a specific protein target. Instead of simply putting attention on the diversity of molecule structure, we explore the possibility of generating novel molecules based on the existing knowledge. Here, we designed a deep reinforcement learning reward (DRL-reward) function that consists of the final property score, containing specific fragments (CSF) score and pharmacophore score as:

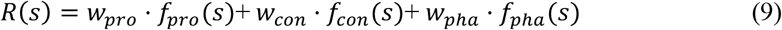

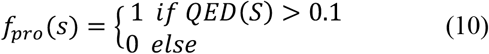

where *w*_*pro*_ represents weight for the quantitative estimate of drug-likeness (QED) property and its default value is 0.1. *f*_*pro*_ stands for the QED. QED values can range from 0 (all properties are unfavourable) to 1 (all properties are favourable), which are calculated by eight molecular properties.^53^ The score function *f*_*csf*_ of containing specific fragments (csf) is:

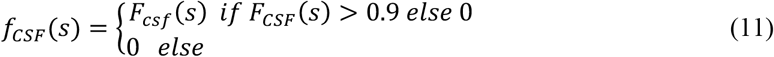

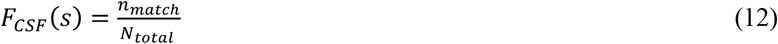

The binding site of SARS-CoV-2 3CL^pro^ (Figure 9) is commonly divided into the catalytic activity center (His41 and Cys145, specified as S1’) and several subsites, defined as S1 (His163, Glu166, Phe140, Leu141 and Asn142), S1’ (His41, Cys145, Gly143 and Ser144), S2 (Tyr54, Asp187, His41, Arg188, His164 and Met49), and S3 (Thr190, Gln192, Glu166, Met165, Leu167 and Gln189). Each subsite may have its own favorable binding fragment. When generated structures (including the intermediates) containing those favorable fragments, *f*_*csf*_ is equivalent to give an additional reward to our RL agent. *w*_*con*_ controls the contribution of the biased fragment to the reward signal and the default value is 0.6. *n*_*match*_ stands for how many biased fragments have been matched in one generated structure. *N*_*total*_ is the number of biased fragments defined based on our knowledge learned from related work. *f*_*pha*_ represents the score function of pharmacophores, which mainly depends on the ligand-protein interaction mode from the crystal structure (PDB ID: 6LU7):

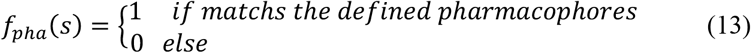

**Figure 9.**
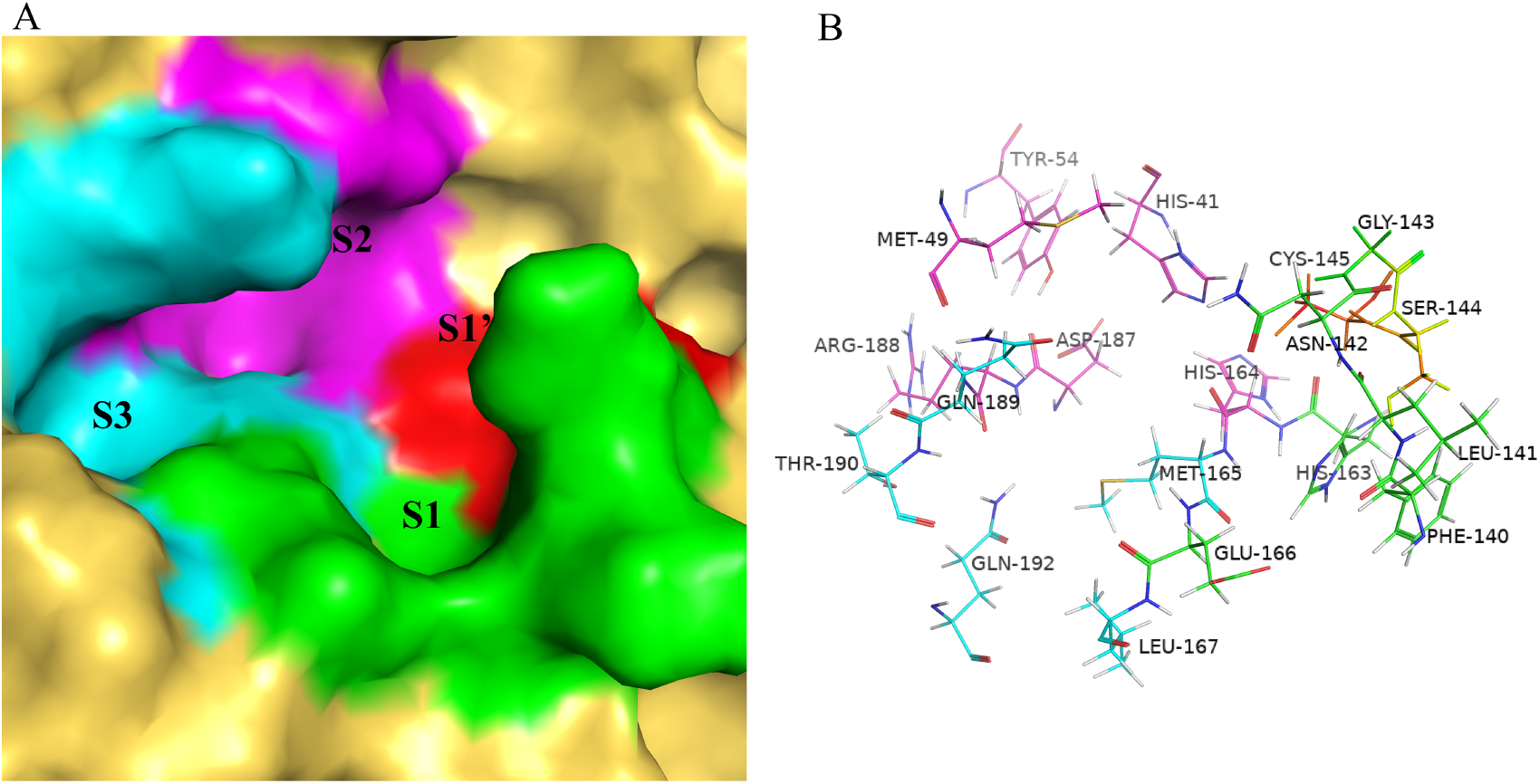
Binding site of SARS-CoV-2 3CL^pro^. A: The subsites that complement the substrate-binding pocket are shown as surface, which are assigned as S1 (green), S1’ (red), S2 (magenta), and S3(cyan). B: The key residues of the binding site are displayed by green, red, magenta and cyan lines (S1, S1’, S2 and S3). The pictures of the binding site are generated using PyMol ((http://www.pymol.org/).

The pharmacophores plot was added in Figure S5. *w*_*pha*_ controls the contribution of the pharmacophore score to the reward and the default value is 0.4.

### Molecular Generation and Selection

As discussed in the above reward design part, our reward function considers the molecular descriptor thresholds (QED>0.1), the defined pharmacophores and biased fragments. In total, 4,922 unique valid structures were automatically generated and all matched the defined rules by using ADQN-FBDD without any pre-training as many other methods need.^35, 40, 41, 54, 55^ Next, All the molecules with high deep reinforcement learning scores (DRL score: R(S)>0.6) were kept (47 molecules). Then, these 47 unique molecules were prepared to generate at least 1 conformation with the local energy minimization using the OPLS-2005 force field by the “ligand prepare” module of Schrödinger 2015 software. The 47 unique molecules generated a total of 163 3D conformations before docking into the substrate-binding site of SARS-CoV-2 3CL^pro^. Considering the balance between precision and calculation time, the stand-precision (SP) Glide^56^ was firstly used to predict the possible non-covalent binding poses in this binding site and the binding site grid centered on the original ligand N3^57^ with 20 Å buffer dimensions. Following this non-covalent docking, we also calculated the covalent docking poses and scores for those 47 molecules. We reordered the docking results mainly based on the covalent docking score and the RMSD value from covalent docking pose to non-covalent (Table S1).

## Supplementary information

Supplementary dataset and additional information of this paper can be found at https://github.com/tbwxmu/2019-nCov.

## Acknowledgements

We would like to thank Prof. Dongqing Wei’s group and the PCL lab for their generous support of high-performance computing resources. BT’s work was funded by the program of China Scholarships Council No. 201806310017. DX’s effort was supported by the US National Institutes of Health grant R35-GM126985.

## Authors’ contributions

BT and DX designed the study. BT developed the AI-aided methods and wrote the manuscript. All authors contributed to the interpretation of results. All authors reviewed and edited the manuscript. All authors read and approved the final manuscript.

## Data and Code availability

The code for the ADQN-FBDD and related data in this paper will be available at https://github.com/tbwxmu/2019-nCov upon acceptance of this paper for journal publication.

## Competing interests

The authors declare that they have no competing interests.

## Notes

https://github.com/tbwxmu/2019-nCov

