## Supplemental Materials for "AI-aided design of novel targeted covalent inhibitors against SARS-CoV-2"

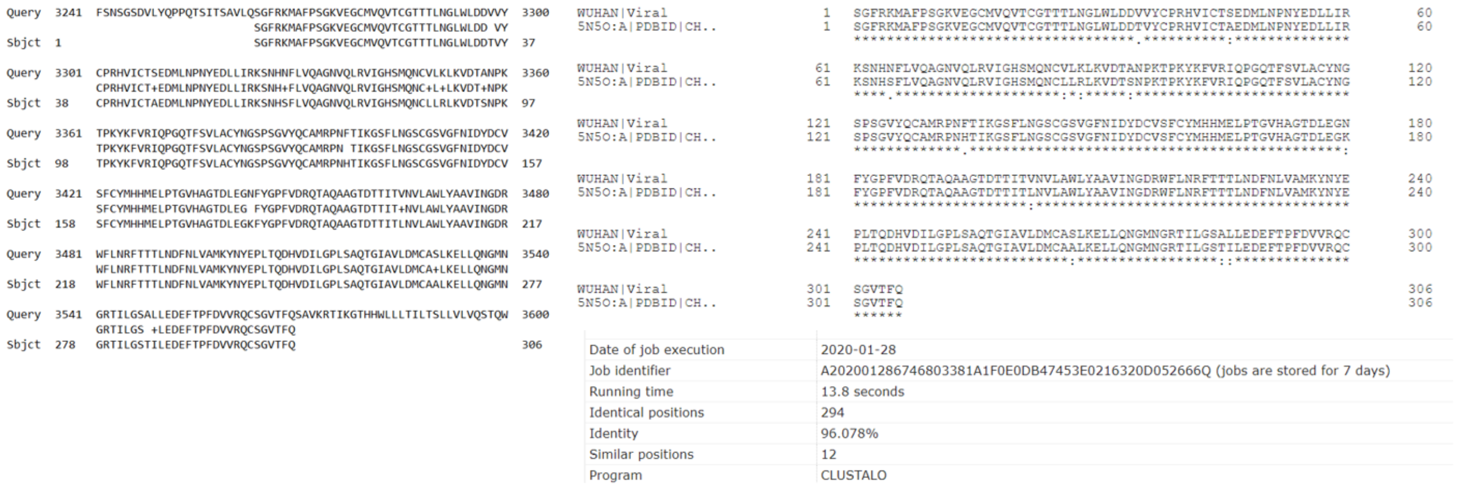


Figure S1. Sequence alignment between SARS-CoV-2 $3CL^{pro}$ and SARS-CoV $3CL^{pro}$.

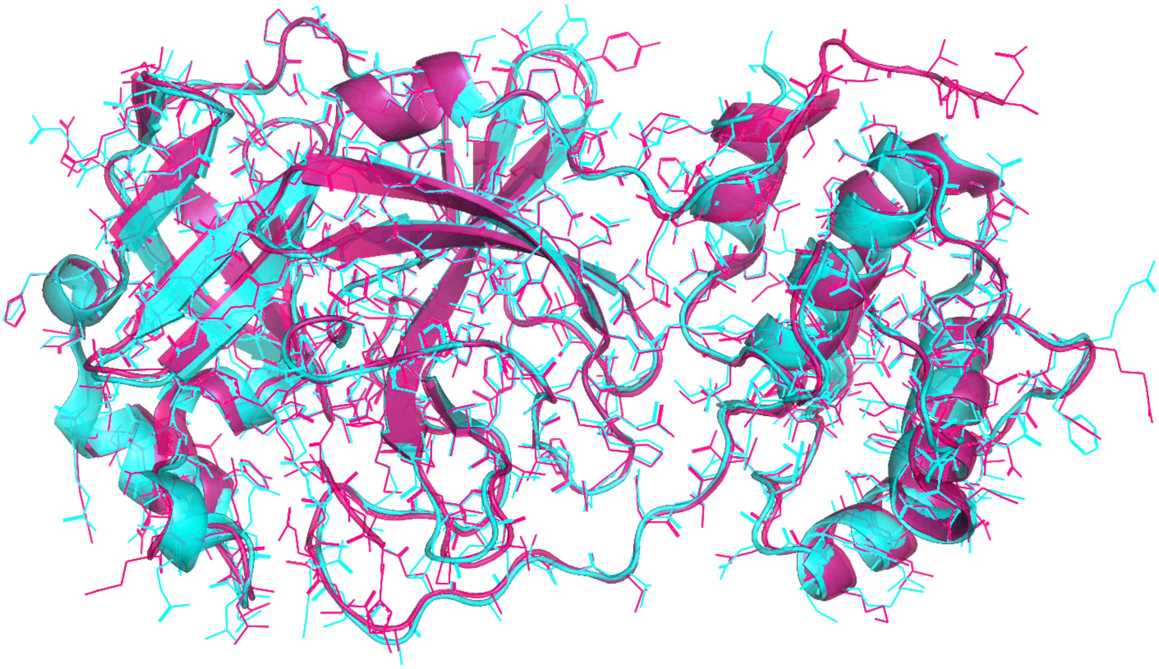

Figure S2. Structure superposition between SARS-CoV-2 3CL$^{pro}$ (PDB ID: 6LU7 shown in magenta) and SARS-CoV 3CL$^{pro}$ (PDB ID: 3D62 shown in cyan) with an RMSD of 0.44 Å.

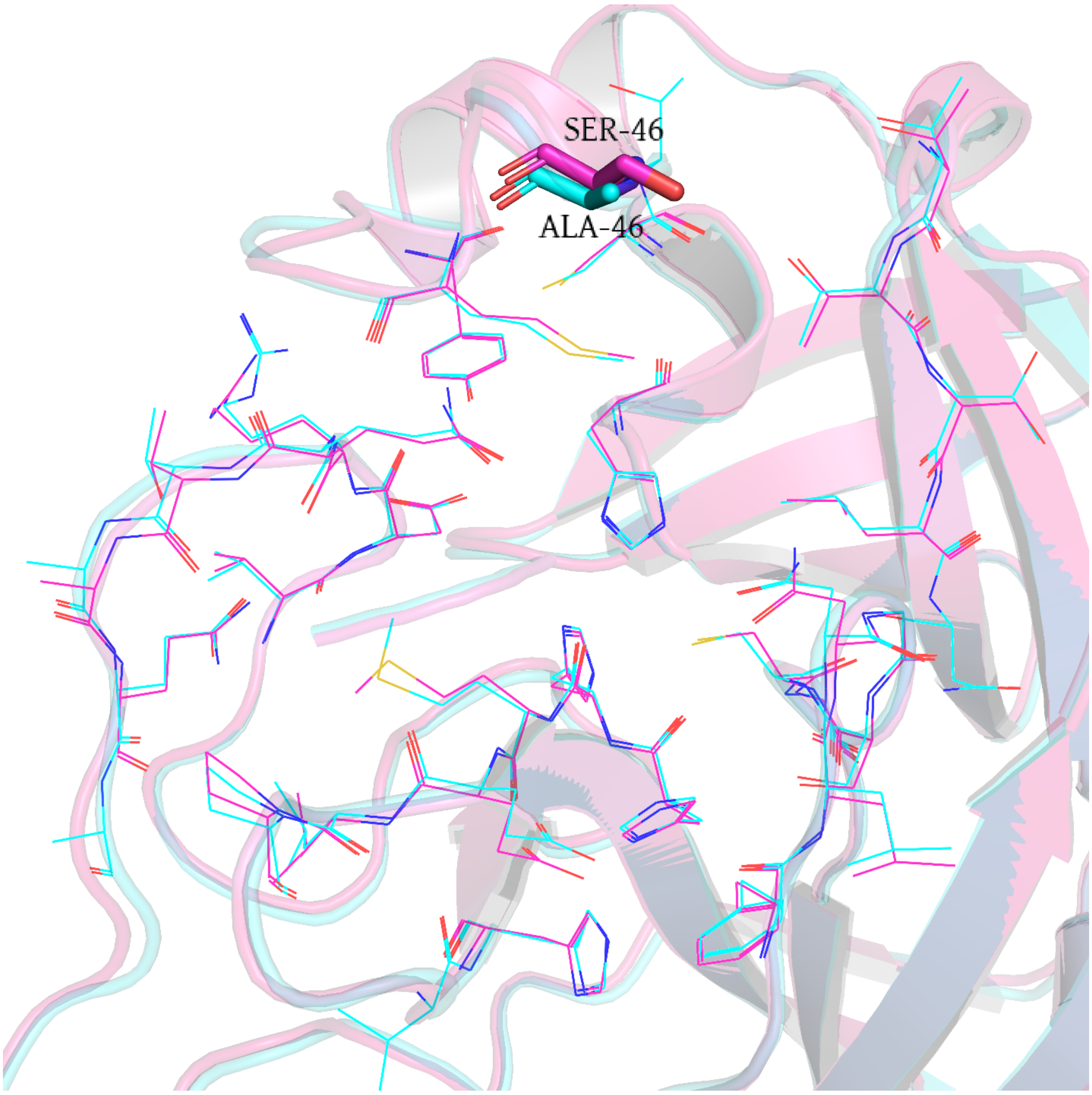


Figure S3. Substrate-binding site superimposition of SARS-CoV-2 3CL$^{pro}$ (PDB ID: 6LU7, magenta cartoon) and SARS-CoV 3CL$^{pro}$ (PDB ID: 2HOB, cyan cartoon). Amino acid residues in the 6Å range of the original molecular ligand N3 (SARS-CoV-2 3CL$^{pro}$: lines in magentas, SARS-CoV 3CL$^{pro}$: lines in cyan) are selected for comparative analysis, and only the residues at position 46 are different. The residue of SARS-CoV-2 3CL$^{pro}$ at position 46 is serine (shown with sticks in magentas), while alanine in SARS-CoV 3CL$^{pro}$ (shown with sticks in cyan).

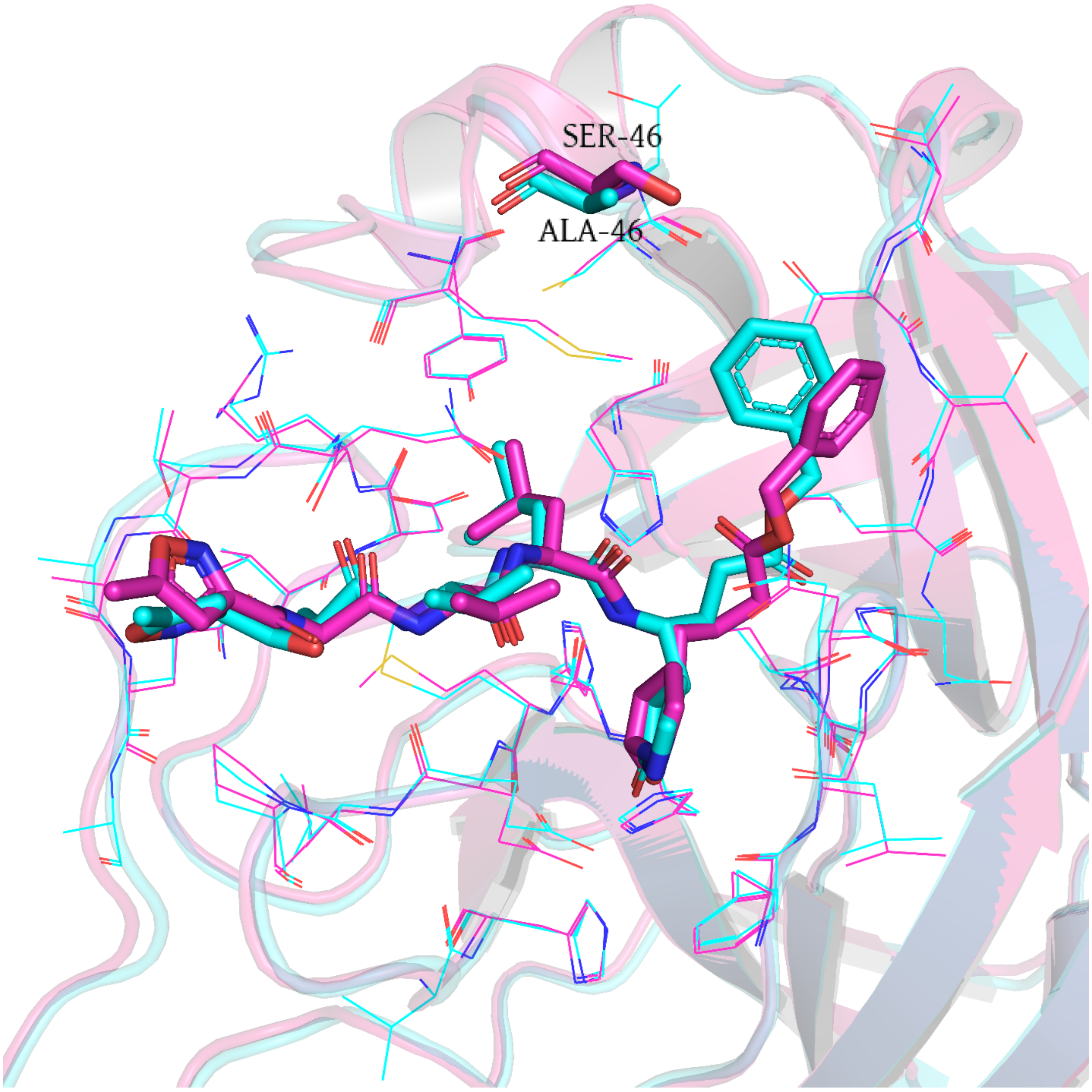


Figure S4. Comparison of the ligand conformations between SARS-CoV-2 3CL$^{pro}$ (PDB ID:6LU7) and SARS-CoV 3CL$^{pro}$ (PDB ID:2HOB). The ligand structure of SARS-CoV-2 3CL$^{pro}$ is shown with sticks in magenta, while the ligand of SARS-CoV 3CL$^{pro}$ is shown with sticks in cyan.

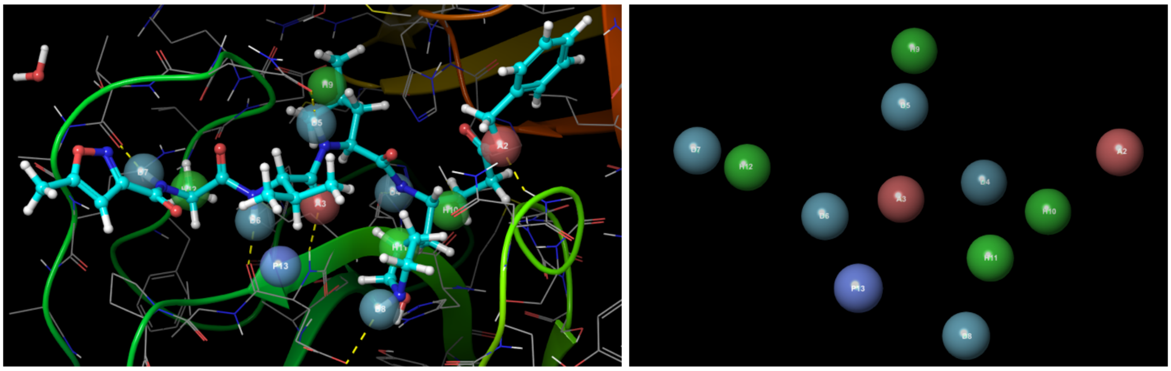

Figure S5. The pharmacophore model embedded in the *R* function. The pharmacophore consists of two hydrogen receptors (red spheres), five hydrogen donors (light blue spheres), four hydrophobic characteristics (green spheres) and a positive charge center (blue spheres).

Tables S1. Results of non-covalent and covalent docking results.

| ID | docking score | cdock affinity | KabschRmsd | StraightRmsd | SMILES |
|---|---|---|---|---|---|
| 1 | -7.63 | | | | N1CC[C@H](C1=O)N[C@H](C(=O)C=O)Cn2nnc(c23)cccc3Oc(c4cccc5)nc(c6c45)cccc6 |
| 2 | -7.743 | -5.464 | 2.752 | 9.245 | N1CC[C@H](C1=O)[C@H](C[C@H]2C)N(CC2)C(=O)/C=C/[C@@](C)(NC(=O)c3ccc(o3)Cl)OCc(c4)ccc(c45)n(C)c(n5)C |
| 3 | -6.802 | -6.374 | 2.381 | 6.455 | c1c[nH]cc(c12)nc(n2)N[C@@](/C=C/C=O)(C[C@@H](C3=O)CCN3)c4c(O)c(c(o4)C=O)-c5c(Cl)cc(F)cc5 |
| 4 | -6.807 | -6.102 | 2.491 | 8.626 | N1CC[C@H](C1=O)C[C@H](N)\C=C\C(=O)Nc(ccc2)c(c2CCC=O)-n3nnc(c34)cccc4C[C@@H](N)CC=O |
| 5 | -8.739 | -8.171 | 4.254 | 8.37 | NC(=O)COC(=O)/C=C/[C@@H](N)C[C@H]1[C@H](Oc(c2cccc3)nc(c4c23)cccc4)OC(=O)[C@@]1([C@@H](C=O)N)C[C@@H](C5=O)CCN5 |
| 6 | -8.772 | -6.262 | 4.314 | 9.295 | FCC(=O)c1ccc(o1)[C@@](N)(C[C@@H](C2=O)CCN2)\C=C\C(=O)N[C@H](C=O)[C@@H](O)Cc(c3)sc(c34)cccc4 |
| 7 | -7.787 | | | | N1CC[C@H](C1=O)N[C@H](C(=O)C=O)Cc2ccc(Cl)cc2 |
| 8 | -7.349 | | | | O=CC(=O)[C@@H](N)Cc1ccnc(c12)c(c(CN)cc2)C[C@@H](C3=O)CCN3 |
| 9 | -8.874 | -7.388 | 1.473 | 1.734 | O=c1c(O)coc(c12)cc(cc2O)NC(=N/[H])\c3c(CCC)cc(cc3)N[C@](N)(/C=C/C=O)C[C@@H](C4=O)CCN4 |
| 10 | -8.967 | -4.159 | 2.393 | 11.36 | N1CC[C@H](C1=O)C[C@@H](N)\C=C\C(=O)N(C)C(=O)[C@H](C)[C@@H](c(s2)ccc2C=O)[C@H](O)[C@@H](N)C(C)(C)c3cccc(c34)cccc4 |
| 11 | -9.075 | -6.923 | 3.952 | 5.132 | N1CC[C@H](C1=O)C[C@@](N)(/C=C/C=O)c2c(cc(s2)C=O)-c(n3)nc(c34)cn(C)cc4N[C@](C=O)(C)c5cnc[nH]5 |
| 12 | -7.256 | -5.275 | 3.352 | 7.935 | O=C\C=C\[C@](C)(N)[C@@H](C1=O)[C@H](OCC=O)CN1C(=O)[C@@H](c(c2)ccc(c23)n(C(=O)C)cc3)C(C)(C)c(c4)sc(c45)cccc5 |
| 13 | -7.545 | -7.222 | 2.834 | 4.748 | N1CC[C@H](C1=O)C[C@](N)(/C=C/C=O)N(C2=O)CCCN2c(cc3O)cc(c34)occ(c4=O)N[C@H](C=O)C |
| 14 | -6.823 | -5.008 | 1.639 | 4.562 | s1cccc1[C@@]([C@H](N)C(=O)C=O)([C@@H](C2=O)CCN2)c3ccc(cc3)\C=C\C=O |
| 15 | -8.301 | | | | N1CC[C@H](C1=O)N[C@H](C(=O)C=O)Cc2c(C3CCC3)cc(F)cc2Cl |
| 16 | -7.663 | -7.199 | 1.837 | 4.789 | N1CC[C@H](C1=O)N[C@H](C(=O)C(=O)O)Cc(c2)sc(c23)cccc3 |
| 17 | -6.766 | -5.437 | 3.614 | 4.207 | O=c(o1)oc(C)c1CN[C@](/C=C/C=O)(C[C@@H](C2=O)CCN2)c3c(ccc(=O)o3)O[C@H]4CO[C@H]([C@H]45)OCC5 |
| 18 | -7.404 | | | | N1CC[C@H](C1=O)C[C@@H](C=O)N(CC=O)[C@H](C(=O)C=O)Cc2cc(N)ccc2 |
| 19 | -7.632 | -8.385 | 2.899 | 5.489 | N1CC[C@H](C1=O)C[C@](N)(/C=C/C=O)[C@H](CC2)C[C@]2(N[C@H](C=O)[C@@H](C)O)c(cc3=O)oc(c34)cc(O)cc4O |
| 20 | -7.234 | -6.597 | 1.873 | 2.615 | N1CC[C@H](C1=O)N[C@H](C(=O)C(=O)O)Cc(c(c2=O)O)oc(c23)c(O)c(O)cc3O |
| 21 | -8.133 | -6.518 | 1.907 | 8.581 | N1CC[C@H](C1=O)C[C@@H](N)\C=C\C(=O)O[C@H]2CO[C@H]([C@H]23)O[C@](C3)(NC(C)(C)CC)N(C(=O)CC)c4ccccc4 |
| 22 | -7.2 | -7.392 | 4.657 | 6.777 | n1nccn1-c(cs2)nc2C(=O)[C@H](C[C@@H](C3=O)CCN3)N[C@H](C=O)[C@H](O)C(=O)N[C@@H](C)\C=C\C=O |
| 23 | -7.625 | -6.972 | 2.61 | 7.52 | c1cccc(c12)c(=O)[nH]n(c2=O)[C@@](N)(\C=C\C(=O)N[C@H](C(=O)O)CC)C[C@@H](C3=O)C[C@@H](N3)C[C@H](N)CC |
| 24 | -7.211 | | | | O=CC(=O)[C@@H](N)Cc(nn1)n(c12)C[C@@](N)(NC2)CCC[C@@H](C3=O)CCN3 |
| 25 | -8.133 | | | | N1CC[C@H](C1=O)N[C@H](C(=O)C=O)Cc2c(O)c(Cl)cc(Cl)c2Cl |
| 26 | -8.346 | -8.322 | 3.402 | 8.209 | C[C@H](N)C(=O)NN(CCC(=O)N)C(=O)c(c(c12)[nH]cc1)c(C[C@@](N)(/C=C/C=O)[C@@H](C3=O)CCN3)cc2C[C@@H](N)CC=O |
| 27 | -7.308 | | | | N1CC[C@H](C1=O)N[C@H](C(=O)C=O)Cc2cccs2 |
| 28 | -6.845 | | | | s1cccc1C[C@@H](C(=O)C=O)NCC[C@@H](N)C[C@@H](C2=O)CCN2 |
| 29 | -8.176 | -7.667 | 1.921 | 2.503 | CC(C)C[C@@H](B(O)O)Nc(cs1)nc1C(=O)[C@H](C)N(C(=O)[C@@H](N)CC)C(=O)/C=C/[C@@H](N)C[C@@H](C2=O)CCN2 |
| 30 | -6.241 | | | | C1CC1C[C@@H](C(=O)C=O)N[C@@H](C2=O)CCN2 |
| 31 | -6.502 | | | | N1CC[C@H](C1=O)N[C@H](C(=O)C=O)CC2CCCC2 |
| 32 | -5.731 | | | | O=CC(=O)[C@@H](N)C[C@@H](C1=O)C[C@@H](C[C@H](N)CC)N1[C@@]2(OCC(C)C)CC[C@@H](O)CC2 |
| 33 | -7.246 | | | | N1CC[C@H](C1=O)[C@H]2N[C@H](C[C@H](N)C(=O)C=O)C[C@@H]([C@@H]23)[C@H](SC)CCC3 |
| 34 | -5.337 | | | | O=CC(=O)[C@@H](N)C[C@@H](C1=O)CCN1 |
| 35 | -5.52 | -5.461 | 2.191 | 6.963 | C[C@H](N)C(=O)C(=O)NC(=O)/C=C/[C@H](N)C[C@@H](C1=O)CCN1 |
| 36 | -4.847 | | | | O=CC(=O)[C@H](C)NC(=O)[C@H](CC(C)C)N[C@@H](C1=O)CCN1 |
| 37 | -7.325 | -5.179 | 1.83 | 2.911 | O=CC(=O)[C@H](C)N(C=O)C(=O)/C=C/[C@H](N)C[C@@H](C1=O)CCN1 |
| 38 | -6.459 | | | | O=CC(=O)[C@@H](N)[C@H](N)[C@@H]1[C@H]([C@@H](C2=O)CCN2)NC[C@@H](N1)N(C)C |
| 39 | -7.998 | -6.302 | 3.796 | 9.2 | n1cc(C)ncc1C(=O)N[C@H](C[C@@H](C2=O)CCN2)\C=C\C(=O)Oc(nc(c3c45)cccc3)c4cc(cc5)-c6ccc([N+]([O-])=O)cc6 |
| 40 | -7.779 | -6.513 | 2.76 | 6.048 | n1ccncc1C(=O)N[C@H](\C=C\C=O)[C@]([C@@H](C2=O)CCN2)(c3ccccc3)Oc(c4cccc5)nc(c6c45)cccc6 |
| 41 | -7.196 | -4.571 | 3.056 | 3.748 | c1ccccc1O[C@@](C)(/C=C/C=O)NC(=O)[C@@H](c2cccnn2)[C@H](C)C[C@@H](C3=O)CCN3c4nccs4 |
| 42 | -7.931 | -4.447 | 4.241 | 6.345 | c1ccccc1\C=C\C(=O)N([C@@H](\C=C\C=O)C[C@@H](C2=O)CCN2)[C@H]([C@@H](O)C[C@H](C=O)C)c3c(F)cc(F)c(F)c3 |
| 43 | -5.985 | | | | O=CC(=O)[C@H](CN)N[C@@H](C1=O)CCN1 |
| 44 | -7.013 | -5.197 | 1.304 | 2.97 | s1cccc1C(=O)N([C@](C)(/C=C/C=O)[C@@H](C2=O)CCN2)c3c(n[nH]n3)Nc(on4)c([N+]([O-])=O)c4C |
| 45 | -8.227 | -5.941 | 3.01 | 6.185 | Cc1onc(C)c1-c(cco2)c2\C=C(\C=O)NC(=O)/C=C/[C@H](C[C@@H](C3=O)CCN3)NC(=O)N[C@H](C=O)C |
| 46 | -8.19 | -8.722 | 1.71 | 2.131 | [nH]1nncc1N([C@@H](C)\C=C\C=O)C(=O)[C@@H]([C@H](C)C[C@@H](C2=O)CCN2)c(cc3=O)oc(c34)cc(O)cc4O |
| 47 | -5.513 | | | | O=CC(=O)[C@H](C)N[C@@H](C1=O)CCN1 |

Several lead compounds do not have the covalent docking results as their reactive group is in an unsuitable direction to 145CYS. All RMSD values are heavy atoms based.

KabschRmsd: Root-mean-square deviation (RMSD) is calculated with the Kabsch algorithm (1976) doi: http://dx.doi.org/10.1107/S0567739476001873).

StraightRmsd: Root-mean-square deviation (RMSD) is calculated directly from the atomic coordinates without superimposing the atoms.

Table S2. 45 rules pf different chemical reactions.

| |
|---|
| [$([C;D3]([#0,#6,#7,#8])(=O)):1]-;!@[$([O;D2]-;!@[#0,#6,#1]):2]>>[1*]-[*:1].[3*]-[*:2] |
| [$([C;D3]([#0,#6,#7,#8])(=O)):1]-;!@[$([N;!D1;!$(N=*);!$(N-[!#6;!#16;!#0;!#1]);!$([N;R]@[C;R]=O)]):2]>>[1*]-[*:1].[5*]-[*:2] |
| [$([C;D3]([#0,#6,#7,#8])(=O)):1]-;!@[$([N;R;$(N(@C(=O))@[C,N,O,S])]):2]>>[1*]-[*:1].[10*]-[*:2] |
| [$([O;D2]-;!@[#0,#6,#1]):1]-;!@[$([C;!D1;!$(C=*)]-;!@[#6]):2]>>[3*]-[*:1].[4*]-[*:2] |
| [$([O;D2]-;!@[#0,#6,#1]):1]-;!@[$([C;$(C(-;@[C,N,O,S])-;@[N,O,S])]):2]>>[3*]-[*:1].[13*]-[*:2] |
| [$([O;D2]-;!@[#0,#6,#1]):1]-;!@[$([c;$(c(:[c,n,o,s]):[n,o,s])]):2]>>[3*]-[*:1].[14*]-[*:2] |

| |
|---|
| [$([O;D2]-;!@[#0,#6,#1]):1]-;!@[$([C;$(C(-;@C)-;@C)]):2]>>[3*]-[*:1].[15*]-[*:2] |
| [$([O;D2]-;!@[#0,#6,#1]):1]-;!@[$([c;$(c(:c):c)]):2]>>[3*]-[*:1].[16*]-[*:2] |
| [$([C;!D1;!$(C=*)]-;!@[#6]):1]-;!@[$([N;!D1;!$(N=*);!$(N-[!#6;!#16;!#0;!#1]);!$([N;R]@[C;R]=O)]):2]>>[4*]-[*:1].[5*]-[*:2] |
| [$([C;!D1;!$(C=*)]-;!@[#6]):1]-;!@[$([S;D2](-;!@[#0,#6])):2]>>[4*]-[*:1].[11*]-[*:2] |
| [$([N;!D1;!$(N=*);!$(N-[!#6;!#16;!#0;!#1]);!$([N;R]@[C;R]=O)]):1]-;!@[$([S;D4]([#6,#0])(=O)(=O)):2]>>[5*]-[*:1].[12*]-[*:2] |
| [$([N;!D1;!$(N=*);!$(N-[!#6;!#16;!#0;!#1]);!$([N;R]@[C;R]=O)]):1]-;!@[$([c;$(c(:[c,n,o,s]):[n,o,s])]):2]>>[5*]-[*:1].[14*]-[*:2] |
| [$([N;!D1;!$(N=*);!$(N-[!#6;!#16;!#0;!#1]);!$([N;R]@[C;R]=O)]):1]-;!@[$([c;$(c(:c):c)]):2]>>[5*]-[*:1].[16*]-[*:2] |
| [$([N;!D1;!$(N=*);!$(N-[!#6;!#16;!#0;!#1]);!$([N;R]@[C;R]=O)]):1]-;!@[$([C;$(C(-;@[C,N,O,S])-;@[N,O,S])]):2]>>[5*]-[*:1].[13*]-[*:2] |
| [$([N;!D1;!$(N=*);!$(N-[!#6;!#16;!#0;!#1]);!$([N;R]@[C;R]=O)]):1]-;!@[$([C;$(C(-;@C)-;@C)]):2]>>[5*]-[*:1].[15*]-[*:2] |
| [$([C;D3;!R](=O)-;!@[#0,#6,#7,#8]):1]-;!@[$([C;$(C(-;@[C,N,O,S])-;@[N,O,S])]):2]>>[6*]-[*:1].[13*]-[*:2] |
| [$([C;D3;!R](=O)-;!@[#0,#6,#7,#8]):1]-;!@[$([c;$(c(:[c,n,o,s]):[n,o,s])]):2]>>[6*]-[*:1].[14*]-[*:2] |
| [$([C;D3;!R](=O)-;!@[#0,#6,#7,#8]):1]-;!@[$([C;$(C(-;@C)-;@C)]):2]>>[6*]-[*:1].[15*]-[*:2] |
| [$([C;D3;!R](=O)-;!@[#0,#6,#7,#8]):1]-;!@[$([c;$(c(:c):c)]):2]>>[6*]-[*:1].[16*]-[*:2] |
| [$([C;!R;!D1;!$(C!-*)]):1]-;!@[$([n;+0;$(n(:[c,n,o,s]):[c,n,o,s])]):2]>>[8*]-[*:1].[9*]-[*:2] |
| [$([C;!R;!D1;!$(C!-*)]):1]-;!@[$([N;R;$(N(@C(=O))@[C,N,O,S])]):2]>>[8*]-[*:1].[10*]-[*:2] |
| [$([C;!R;!D1;!$(C!-*)]):1]-;!@[$([C;$(C(-;@[C,N,O,S])-;@[N,O,S])]):2]>>[8*]-[*:1].[13*]-[*:2] |
| [$([C;!R;!D1;!$(C!-*)]):1]-;!@[$([c;$(c(:[c,n,o,s]):[n,o,s])]):2]>>[8*]-[*:1].[14*]-[*:2] |
| [$([C;!R;!D1;!$(C!-*)]):1]-;!@[$([C;$(C(-;@C)-;@C)]):2]>>[8*]-[*:1].[15*]-[*:2] |
| [$([C;!R;!D1;!$(C!-*)]):1]-;!@[$([c;$(c(:c):c)]):2]>>[8*]-[*:1].[16*]-[*:2] |
| [$([n;+0;$(n(:[c,n,o,s]):[c,n,o,s])]):1]-;!@[$([C;$(C(-;@[C,N,O,S])-;@[N,O,S])]):2]>>[9*]-[*:1].[13*]-[*:2] |
| [$([n;+0;$(n(:[c,n,o,s]):[c,n,o,s])]):1]-;!@[$([c;$(c(:[c,n,o,s]):[n,o,s])]):2]>>[9*]-[*:1].[14*]-[*:2] |
| [$([n;+0;$(n(:[c,n,o,s]):[c,n,o,s])]):1]-;!@[$([C;$(C(-;@C)-;@C)]):2]>>[9*]-[*:1].[15*]-[*:2] |
| [$([n;+0;$(n(:[c,n,o,s]):[c,n,o,s])]):1]-;!@[$([c;$(c(:c):c)]):2]>>[9*]-[*:1].[16*]-[*:2] |
| [$([N;R;$(N(@C(=O))@[C,N,O,S])]):1]-;!@[$([C;$(C(-;@[C,N,O,S])-;@[N,O,S])]):2]>>[10*]-[*:1].[13*]-[*:2] |
| [$([N;R;$(N(@C(=O))@[C,N,O,S])]):1]-;!@[$([c;$(c(:[c,n,o,s]):[n,o,s])]):2]>>[10*]-[*:1].[14*]-[*:2] |
| [$([N;R;$(N(@C(=O))@[C,N,O,S])]):1]-;!@[$([C;$(C(-;@C)-;@C)]):2]>>[10*]-[*:1].[15*]-[*:2] |
| [$([N;R;$(N(@C(=O))@[C,N,O,S])]):1]-;!@[$([c;$(c(:c):c)]):2]>>[10*]-[*:1].[16*]-[*:2] |
| [$([S;D2](-;!@[#0,#6])):1]-;!@[$([C;$(C(-;@[C,N,O,S])-;@[N,O,S])]):2]>>[11*]-[*:1].[13*]-[*:2] |
| [$([S;D2](-;!@[#0,#6])):1]-;!@[$([c;$(c(:[c,n,o,s]):[n,o,s])]):2]>>[11*]-[*:1].[14*]-[*:2] |
| [$([S;D2](-;!@[#0,#6])):1]-;!@[$([C;$(C(-;@C)-;@C)]):2]>>[11*]-[*:1].[15*]-[*:2] |
| [$([S;D2](-;!@[#0,#6])):1]-;!@[$([c;$(c(:c):c)]):2]>>[11*]-[*:1].[16*]-[*:2] |
| [$([C;$(C(-;@[C,N,O,S])-;@[N,O,S])]):1]-;!@[$([c;$(c(:[c,n,o,s]):[n,o,s])]):2]>>[13*]-[*:1].[14*]-[*:2] |
| [$([C;$(C(-;@[C,N,O,S])-;@[N,O,S])]):1]-;!@[$([C;$(C(-;@C)-;@C)]):2]>>[13*]-[*:1].[15*]-[*:2] |
| [$([C;$(C(-;@[C,N,O,S])-;@[N,O,S])]):1]-;!@[$([c;$(c(:c):c)]):2]>>[13*]-[*:1].[16*]-[*:2] |
| [$([c;$(c(:[c,n,o,s]):[n,o,s])]):1]-;!@[$([c;$(c(:[c,n,o,s]):[n,o,s])]):2]>>[14*]-[*:1].[14*]-[*:2] |
| [$([c;$(c(:[c,n,o,s]):[n,o,s])]):1]-;!@[$([C;$(C(-;@C)-;@C)]):2]>>[14*]-[*:1].[15*]-[*:2] |
| [$([c;$(c(:[c,n,o,s]):[n,o,s])]):1]-;!@[$([c;$(c(:c):c)]):2]>>[14*]-[*:1].[16*]-[*:2] |
| [$([C;$(C(-;@C)-;@C)]):1]-;!@[$([c;$(c(:c):c)]):2]>>[15*]-[*:1].[16*]-[*:2] |
| [$([c;$(c(:c):c)]):1]-;!@[$([c;$(c(:c):c)]):2]>>[16*]-[*:1].[16*]-[*:2] |

Note, a reaction definition is based on the SMILES arbitrary target specification (SMARTS), which is a chemical language for describing molecular patterns. We also use the same reaction type for adding fragments, except with the reverse reaction.
